# A meta-analysis reveals temperature, dose, life stage, and taxonomy influence host susceptibility to a fungal parasite

**DOI:** 10.1101/818377

**Authors:** Erin L. Sauer, Jeremy M. Cohen, Marc J. Lajeunesse, Taegan A. McMahon, David J. Civitello, Sarah A. Knutie, Karena Nguyen, Elizabeth A. Roznik, Brittany F. Sears, Scott Bessler, Bryan K. Delius, Neal Halstead, Nicole Ortega, Matthew D. Venesky, Suzanne Young, Jason R. Rohr

**Affiliations:** Department of Integrative Biology, University of South Florida, Tampa, FL; Department of Forest and Wildlife Ecology, University of Wisconsin, Madison, WI; Department of Biology, University of Tampa, Tampa, FL; Department of Biology, Emory University, Atlanta, GA; Department of Ecology and Evolutionary Biology, University of Connecticut, Storrs, CT; Department of Research and Conservation, Memphis Zoo, Memphis, TN; BioScience Writers, Houston, TX; Wildlands Conservation, Tampa, FL; Department of Biology, Allegheny College, Meadville, PA; Environmental Engineering Institute, Ecole polytechnique fédérale de Lausanne (EPFL), Lausanne, Switzerland; Department of Biological Science, University of Notre Dame, South Bend, IN

**Keywords:** Meta-analysis, *Batrachochytrium dendrobatidis*, amphibian declines, experimental design, disease ecology, chytridiomycosis, thermal mismatches, dose response, life stage effects

## Abstract

Complex ecological relationships, such as host-parasite interactions, are often modeled with laboratory experiments. However, some experimental laboratory conditions, such as temperature or infection dose, are regularly chosen based on convenience or convention and it is unclear how these decisions systematically affect experimental outcomes. Here, we conducted a meta-analysis of 58 laboratory studies that exposed amphibians to the pathogenic fungus *Batrachochytrium dendrobatidis* (Bd) to better understand how laboratory temperature, host life stage, infection dose, and host species affect host mortality. We found that host mortality was driven by thermal mismatches: hosts native to cooler environments experienced greater Bd-induced mortality at relatively warm experimental temperatures and vice versa. We also found that Bd dose positively predicted Bd-induced host mortality and that the superfamilies Bufonoidea and Hyloidea were especially susceptible to Bd. Finally, the effect of Bd on host mortality varied across host life stages, with larval amphibians experiencing lower risk of Bd-induced mortality than adults or metamorphs. Metamorphs were especially susceptible and experienced mortality when inoculated with much smaller Bd doses than the average dose used by researchers. Our results suggest that when designing experiments on species interactions, researchers should carefully consider the experimental temperature, and inoculum dose, and life stage and taxonomy of the host species.

## Introduction

Laboratory experiments are a common tool used in ecology to better understand complex species interactions. However, experimental laboratory conditions are often chosen based on convenience or convention, which could intentionally or unintentionally affect experimental outcomes (Hairston 1989). For example, experiments on temperature-dependent organisms are often conducted at a constant temperature with no justification as to why that temperature was chosen. Furthermore, researchers might select convenient or conventional populations, strains, and densities of organisms to study (Hairston 1989). The consequences of these common decisions for the outcomes of species interaction studies are often not well understood. However, methodological choices are likely to bias extrapolations of experimental results to the population level under natural conditions, potentially affecting management decisions. Disentangling these potential confounding effects of experimental design on host-parasite interactions is therefore critical, especially because of the recent rise in emerging infectious diseases that are causing global declines in biodiversity (Goulson et al. 2015, Scheele et al. 2019). Thus, researchers must understand how test conditions intentionally or unintentionally alter experimental outcomes and affect host-parasite interactions and disease progression, especially for systems in which experimental outcomes can inform resource management and conservation.

The pathogenic fungus *Batrachochytrium dendrobatidis* (Bd) has been associated with hundreds of amphibian declines worldwide over the past 40 years (Scheele et al. 2019). Consequently, Bd has been the focus of thousands of studies and surveys in recent decades, many of which have been used to inform conservation efforts (Skerratt et al. 2007, Rohr and Raffel 2010, Converse et al. 2017). Bd infects keratinized mouthparts of larval amphibians and keratinized skin on the whole body of post-metamorphic amphibians, degrading the epithelial layer and causing the lethal disease chytridiomycosis (Berger et al. 1998, Pessier et al. 1999, Grogan et al. 2018). Effects of the pathogen on wild host populations vary greatly, with some species experiencing declines, extirpations, or even total extinction, and others experiencing few to no negative impacts (Venesky et al. 2014, Berger et al. 2016, Scheele et al. 2019). Heterogeneity in virulence, tolerance, and resistance among species has led to conflicting findings and much debate about mechanisms that might be driving these apparent differences in mortality risk among host populations (Fisher et al. 2009).

Many factors affect Bd-host interactions (Blaustein et al. 2018), including host behavior (Sauer et al. 2018), body size (Carey et al. 2006), Bd isolate (O’Hanlon et al. 2018), zoospore dose (Carey et al. 2006), temperature (Cohen et al. 2017), host taxon (Gervasi et al. 2017), and life stage (McMahon and Rohr 2015). However, the influences of these factors on Bd-host interactions are often not straightforward and not well-understood. For example, many studies have independently concluded that warm temperatures are positively associated with Bd prevalence and host mortality (e.g., Pounds et al. 2006, Bosch et al. 2007), while many others have found associations between Bd outbreaks and cold temperatures or seasons (e.g., Retallick et al. 2004, Kriger and Hero 2007). Recently, a more context-dependent hypothesis, the thermal mismatch hypothesis, was proposed to explain these inconsistencies. This hypothesis suggests that host species adapted to warmer climates should be more susceptible to disease at relatively cool temperatures, whereas cool-adapted host species should be most susceptible during unusually warm periods (Appendix S1: Fig. S1, Cohen et al. 2017). This hypothesis assumes that smaller-bodied pathogens generally have wider thermal breadths than their larger-bodied hosts (Rohr et al. 2018) and are limited by extremes, which allows pathogens to outperform their hosts under abnormal, but not extreme conditions (Cohen et al. 2017). The thermal mismatch hypothesis is supported by multiple laboratory experiments and global-scale analyses of data collected in the field which show warm- and cool-adapted hosts experience faster Bd growth and greater Bd prevalence at cool and warm temperatures, respectively (Cohen et al. 2017, Sauer et al. 2018, Cohen et al. 2019a, Cohen et al. 2019b). However, experimental evidence for this hypothesis is restricted to only three amphibian host species (Cohen et al. 2017, Sauer et al. 2018).

Conversely, more is understood about the effects of life stage, zoospore dose, and host taxa on Bd-amphibian interactions than the effects of temperature, but the generality of these effects has not been explored in detail. For example, dose (number of infective Bd zoospores a host is exposed to) typically increases host mortality risk (Carey et al. 2006). However, it is unclear which doses are generally needed to produce mortality across life stages, information that would be useful for researchers examining sub-lethal effects of Bd. Finally, many researchers select local, easily-collected study species for experiments out of convenience, but species vary greatly in their susceptibility to Bd (Gervasi et al. 2017). Field evidence has suggested that globally, tropical Bufonidae and Hyloidea species have undergone more severe declines than other groups of amphibians (Scheele et al. 2019). However, mortality risk does not always translate to extinction risk. For example, a cross-taxon study of twenty Anuran species in North America found that Bufonidae species were more susceptible to Bd than Ranidae or Hylidae species despite a greater number of documented Bd-associated declines among Ranidae and Hylidae in North America (Gervasi et al. 2017, Scheele et al. 2019). Thus, a global synthesis of existing data is needed to determine which amphibian taxa have the greatest risk of mortality following Bd exposure independent of factors that might increase extinction risk in the wild (e.g. local climate or restricted range sizes).

Understanding how different experimental factors might unexpectedly affect Bd-induced amphibian mortality would allow researchers to better design experiments, target the intended research question, and appropriately apply experimental findings to conservation efforts. Here, we use a meta-analysis of 58 experimental studies conducted in the amphibian-Bd system to assess how common experimental factors affect Bd-induced mortality risk in amphibian hosts. First, we asked if amphibian host susceptibility to Bd is dependent on temperature, predicting that mortality increases when there is a greater mismatch between laboratory temperature and the mean temperature to which the host is adapted. Second, we examined how host life stage (larva, metamorph, or adult) influences susceptibility to Bd, expecting that metamorphs and adults would be more susceptible to Bd than larvae because more of their skin is keratinized. Third, we asked whether Bd dose affects host mortality, predicting that mortality risk increases with Bd dose and that metamorphs are at greatest risk of Bd-induced mortality at relatively low doses. Finally, we examined how host taxonomy influences susceptibility to Bd, expecting that Bufonidae and Hyloidea species would be most susceptible to Bd given the severe Bd-associated declines in those groups. To accomplish these four goals, we searched the published literature for amphibian-Bd laboratory studies and modeled effects of thermal mismatches, life stage, and dose on Bd-induced mortality.

## Materials and Methods

### Data collection

Our goal was to synthesize all experimental studies that compared amphibian hosts experimentally infected with Bd in the laboratory to unexposed controls. To accomplish this, we conducted a meta-analysis, which allowed us to standardize and combine results across multiple experiments to draw a broader conclusion than could be typically drawn from any one experiment. We located studies in Web of Science by searching for the term “*Batrachochytrium dendrobatidis”* in October 2016, producing 1,403 results. We included laboratory studies meeting all of the following conditions: 1) at least one Bd-exposed treatment paired with an unexposed control group (we did not consider treatments that exposed hosts to additional parasites [e.g. co-infection] or pesticides and other compounds), 2) treatments held at a constant laboratory temperature, 3) hosts were either wild-collected or lab-reared from wild-collected parents to avoid situations where hosts may have adapted to the climatic conditions of the captive-breeding facility, 4) treatment and control mortality, sample sizes, and host and Bd isolate collection location were either available in the manuscript or provided to us by the author when requested (final count: 58 studies). See Appendix S1 for more details regarding data collection.

### Effect sizes

In meta-analyses, effect sizes must be calculated to provide a standardized measure of an effect across studies (Borenstein et al. 2011). Because mortality data are binary, we calculated log odds ratios to assess the odds of mortality in the Bd-exposed animals relative to the control animals (Cox 2018), using the following equation:

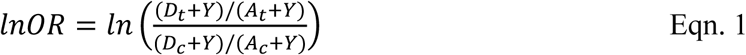

where *D*_*t*_ is the number of treatment (i.e. Bd-exposed) animals that died, *A*_*t*_ is the number of treatment animals that survived, *D*_*c*_ is the number of control (i.e. sham-exposed) animals that died, *A*_*c*_ is the number of control animals that survived, and *Y* (*Y* = 1/2) is a Yate’s continuity correction to avoid error in our effect sizes resulting from dividing by zero (Yates 1934). Yate’s continuity correction (*Y*) was only added to effect sizes and variance equations where an error from dividing by zero would have occurred; all other effect sizes and variances were calculated using the same log odds ratio formula but with *Y* omitted (Sweeting et al. 2004). When conducting analyses, odds ratios must be natural log-transformed to ensure that studies with equal but opposite effects have odds ratios that differ from zero by the same magnitude but in opposite directions (Borenstein et al. 2011). Variance for each effect size was calculated as:

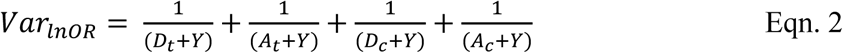

A log odds ratio significantly greater than zero represents greater mortality in the treatment group than in the control, whereas a log odds ratio with 95% confidence intervals that overlap with zero represents a failure to reject the null hypothesis that Bd exposure has no effect on host survival. We calculated log odds ratios from mortality reported at the end of each experiment, regardless of experimental duration. However, mortality tends to increase over time and studies varied in their duration. We were unable to conduct a time series analysis without losing a large portion of studies, as many did not report survival over time. Therefore, we controlled for inconsistencies in experimental length by including duration of experiment as a moderator in our model (see *Statistical analysis* section).

### Statistical analysis

All analyses were conducted in R 3.5.1 (2017). We analyzed the data using a mixed-effects meta-analysis (*metafor* package, *rma.mv* function (2010)), described with the following regression equation:

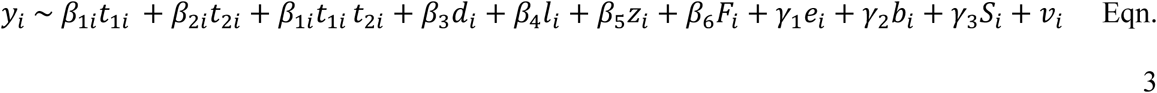

Where ***y***_***i***_ denotes log odds ratios and ***v***_***i***_ denotes log odds ratio variance for the *i*^th^ effect size. Our primary hypotheses concerned the relationship between experimental conditions and Bd infection outcome, not simply the main effect of Bd on host mortality. Therefore, our models included the following multiple moderators. First, thermal mismatch effect (***t***_***1***_ * ***t***_***2***_), was represented by an interaction between 50-year mean temperature at the host’s collection site extracted from WorldClim (***t***_***1***_; assumed host-adapted temperature) and the laboratory temperature (***t***_***2***_) at which the experiment was conducted. We also considered using mean annual minimum and maximum temperatures as the expected temperature to which hosts have adapted and have included those analyses as well as further explanation for our use of long-term annual mean temperature as assumed host-adapted temperature in Appendix S1 (see Appendix S1: Data collection & Figure S1). Support for the thermal mismatch hypothesis is represented by a negative interaction between these two factors, where cool- and warm-adapted hosts experience the greatest Bd-induced mortality at warm and cool laboratory temperatures, respectively. We also included moderators for effects of experimental duration (***d***), life stage (***l***; three-level categorical variable: larvae, metamorph, adult), log_10_-transformed Bd zoospore dose (***z***), and taxonomic group (***F***; six-level categorical variable). In order to explore differences in susceptibility among host taxa, species were consolidated into taxonomic groups with larger sample sizes. Thus, taxonomic groups represent either a superfamily (Bufonoidea, Hyloidea, Ranoidea, and Pelobatoidea), or a suborder (Salamandroidea and Archaeobatrachia). See Appendix S1 and Appendix S1: Table S1 for summary information and full list of host species included in the meta-analysis.

To avoid bias and risk of type I error, we accounted for between-study random effects (***e***) as well as non-independence among Bd isolates by including Bd isolate (***b***) and host species (***S***) as a random intercept in our models (Borenstein et al. 2011, Civitello et al. 2015). Due to the complex non-independence among effect sizes within a study (e.g. some studies had multiple effect sizes), we did not use funnel plots or rank correlation tests to assess publication bias (Lau et al. 2006, Civitello et al. 2015). To create the partial residual plots, which allowed us to visualize the main effects and interactions in our model (Figs. 2, 3, & 4) while controlling for other covariates in the model, we created an identical meta-analytic model using a Bayesian linear mixed-effects package (*blme* package, *blmer* function (2013); see Appendix S1 for more details) then generated plots using the *visreg* package (Breheny and Burchett 2013). The coefficients and error estimates generated from the *blme* model were identical to the results generated by the *metafor* model (see Appendix S1: Table S2 for comparison). We used this approach because visualization tools for mixed-effects meta-analytic models in *metafor* are currently limited. We report summary statistics and *p*-values from our *metafor* model summary because *metafor* is explicitly intended to be used for meta-analysis and thus reports the appropriate summary statistics, p-values, and confidence intervals while *blme* does not.

**Figure 1.**
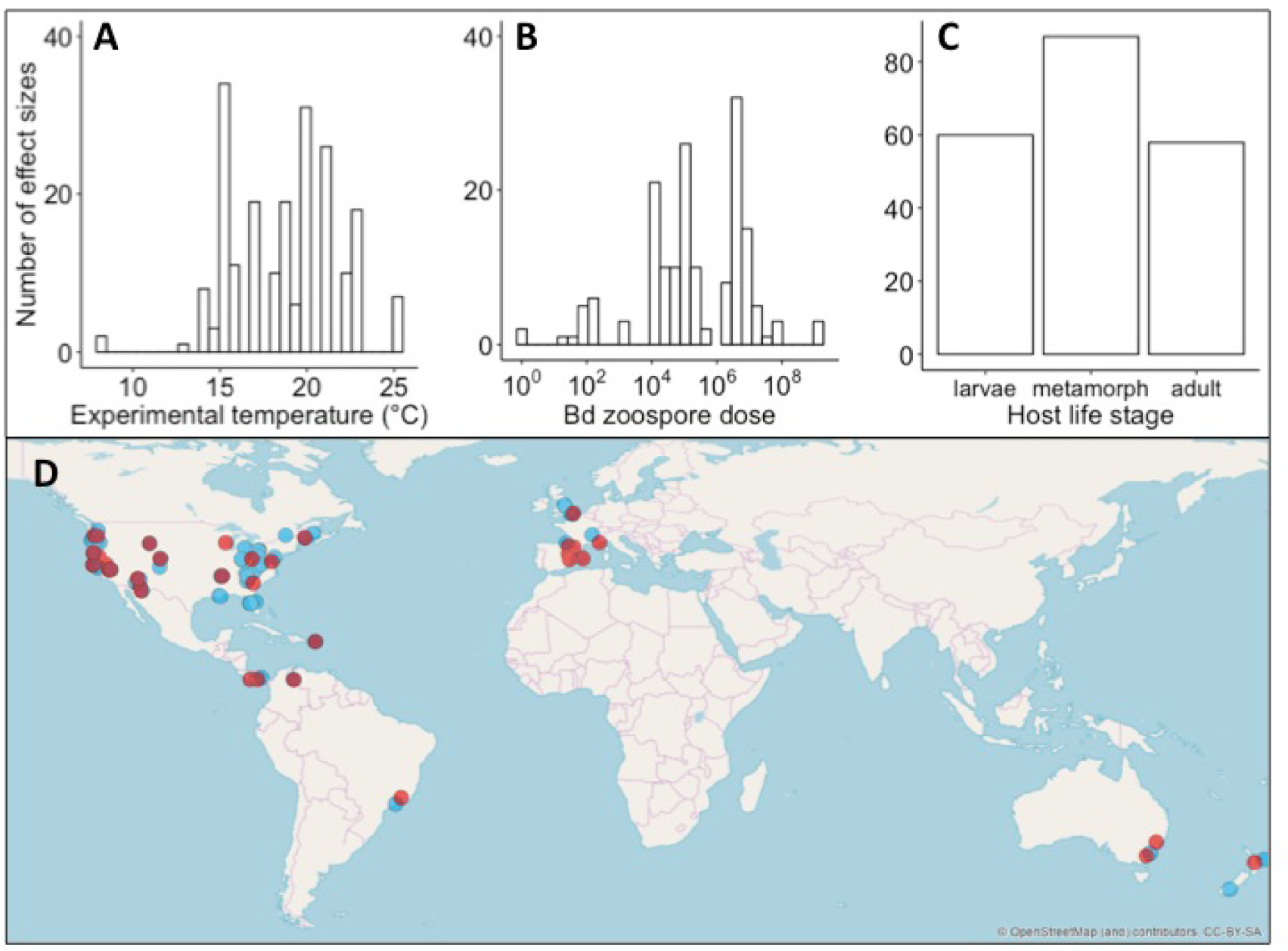
Distribution of (A) experimental temperatures, (B) Bd zoospore doses, and (C) host life stages used in Bd-amphibian experiments for studies included in the meta-analysis. (D) Map showing where hosts (blue points) and Bd isolates (red points) were collected from for studies included in the meta-analysis. All points on the map have the same opacity; locations where points appear darker indicate spatial overlap.

**Figure 2.**
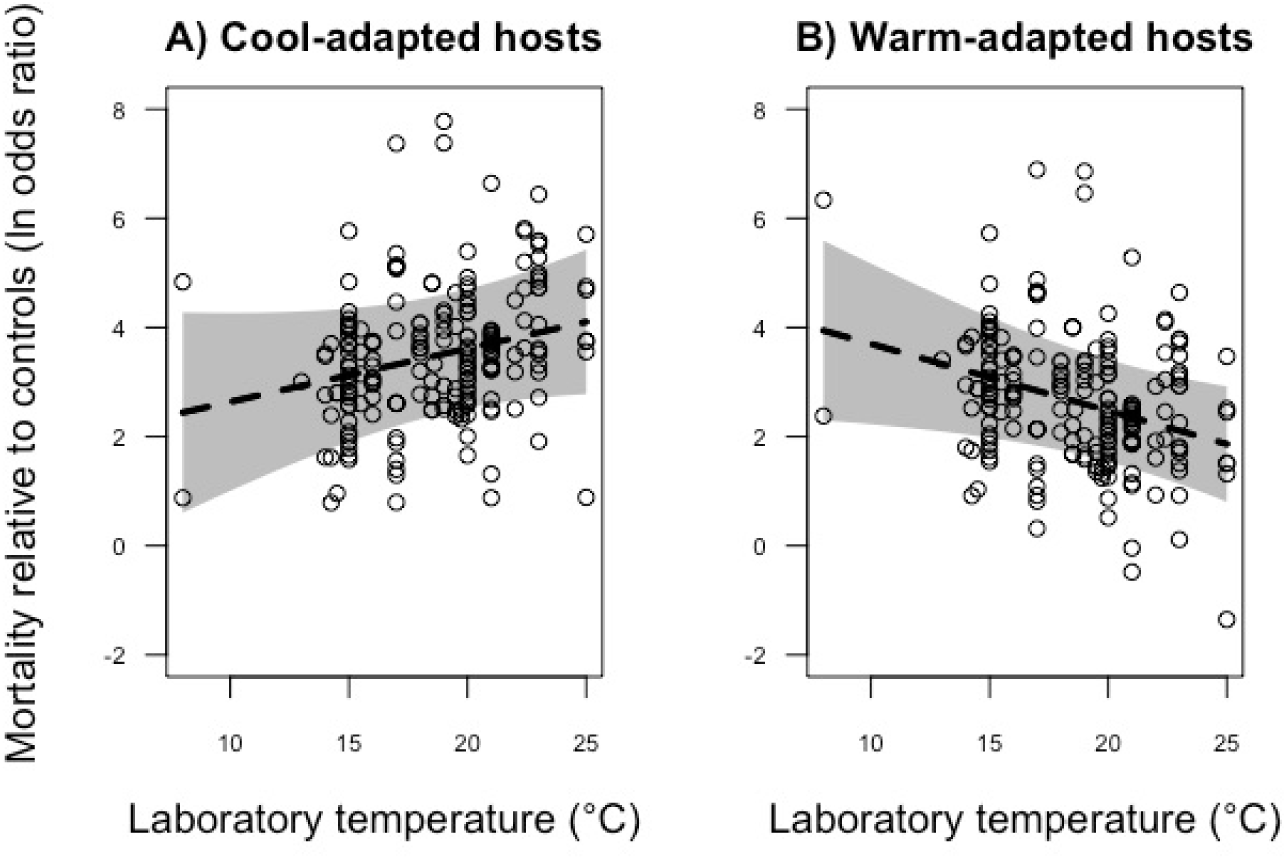
Partial residual plot showing the predicted effect of laboratory temperature on mortality of (A) cold-adapted (20th percentile 50-year mean temperature, or climate: 4.8 °C) and (B) warm-adapted hosts (80th percentile 50-year mean temperature: 16.2 °C) relative to control animals (presented as an odds ratio on the y-axis) between 29 and 42 days after Bd exposure. The plot displays the significant two-way interaction between historic 50-year mean temperature at the collection location of the host and experimental laboratory temperature. Positive values indicate greater mortality among Bd-exposed animals than among unexposed control animals. Points represent individual studies included in the meta-analysis and grey-shading shows associated 95% credible bands.

**Figure 3.**
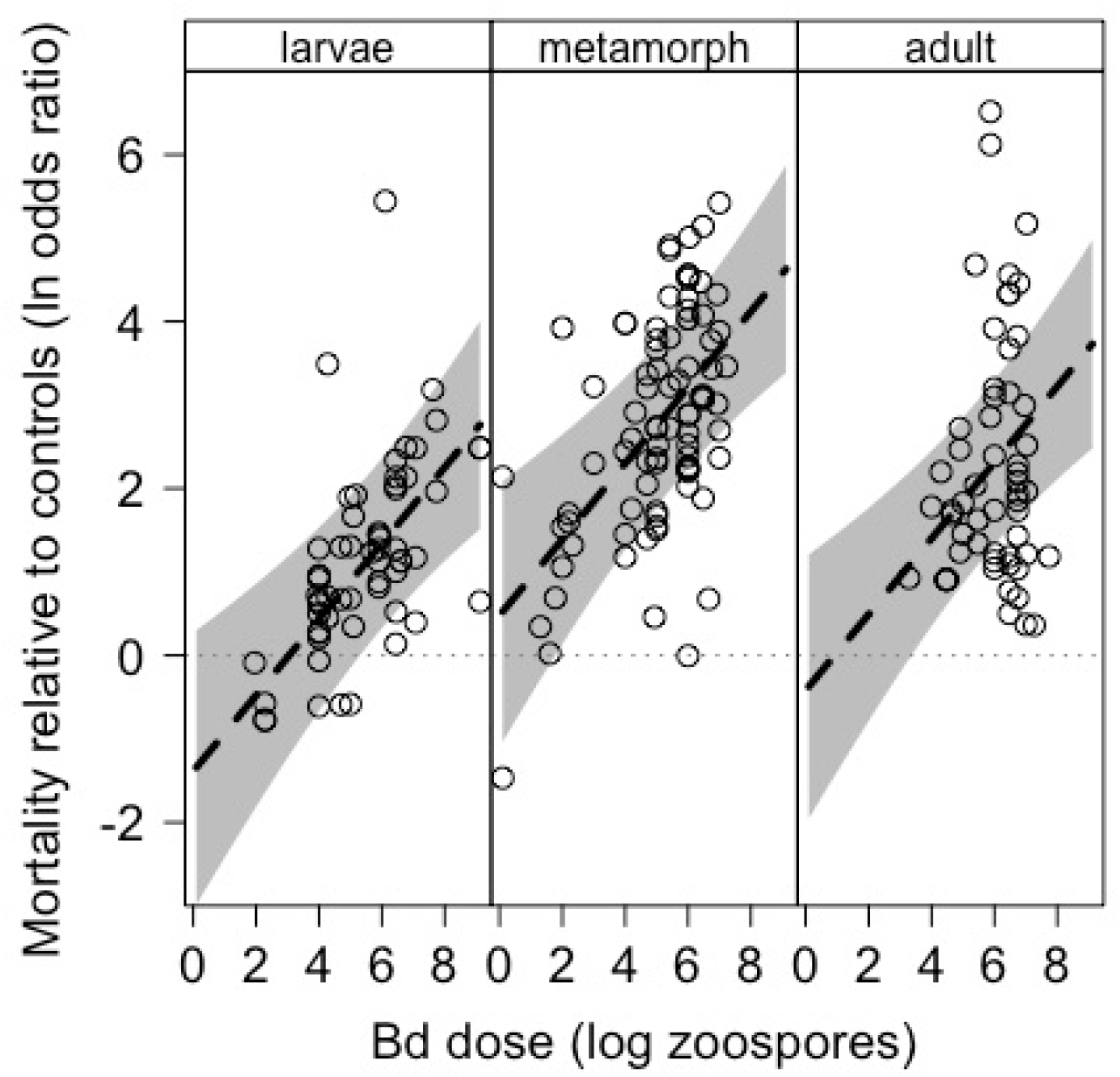
Partial residual plot showing the effect of Bd dose (log_10_(zoospores)) on mortality of Bd-exposed animals relative to controls (presented as natural log odds ratio) on A) larvae, B) metamorphs, and C) adult hosts. Positive values indicate greater mortality among Bd-exposed animals than among unexposed control animals. Points represent individual studies included in the meta-analysis and grey-shading shows associated 95% credible bands.

## Results

Our literature search yielded 205 effect sizes from 58 studies and included 47 amphibian species from 11 families. Experiments used a total of 45 unique Bd isolates (DataS1: Database S1). Host species were collected from North and South America, Europe, and Oceania (Fig. 1). Surprisingly, there were no studies that met our inclusion criteria from the Middle East, Asia, or Africa (Fig. 1).

When controlling for among-study variance, Bd isolate, host species, and experimental duration, we found a negative interaction between host-adapted temperature and laboratory temperature (thermal mismatch effect) (z = −2.75 *p* < 0.01; Table 1 & Fig. 2); cool-adapted hosts experienced the greatest mortality relative to controls at warm laboratory temperatures and warm-adapted hosts experienced the highest mortality relative to controls at cool laboratory temperatures (Table 1 & Fig. 2).

**Table 1.** Results of a mixed-effects meta-analytic model of experiments relating mortality caused by *Batrachochytrium dendrobatidis* to factors including, temperature, life stage, dosage, and taxon. The main effect of Bd on mortality is indicated by the grand mean whereas the coefficient represents the pooled effect of Bd on mortality as a log odds ratio. For continuous variables, asterisks indicate a slope that deviates significantly from zero where the sign of the coefficient indicates the direction of the effect. For categorical variables (life stage and host taxa), asterisks indicate a significant difference from the grand mean with the coefficient indicating the direction and the difference from the grand mean.

|  | Coefficient | SE | z value | p value |  |
| --- | --- | --- | --- | --- | --- |
| Grand mean | 1.559 | 0.314 | 4.964 | < 0.001 | * |
| LongTermTemp | 0.288 | 0.133 | 2.164 | 0.031 | * |
| LabTemp | 0.189 | 0.094 | 1.999 | 0.046 | * |
| LongTermTemp*LabTemp | -0.019 | 0.007 | -2.746 | 0.006 | * |
| Duration | 0.009 | 0.004 | 2.159 | 0.031 | * |
| Larvae | -0.947 | 0.228 | -3.983 | < 0.001 | * |
| Metamorph | 0.924 | 0.195 | 4.740 | < 0.001 | * |
| Adult | 0.023 | 0.266 | 0.087 | 0.931 |  |
| logDose | 0.454 | 0.114 | 3.997 | < 0.001 | * |
| Bufonoidea | 1.690 | 0.657 | 2.573 | 0.010 | * |
| Hyloidea | 1.093 | 0.601 | 1.817 | 0.069 |  |
| Ranoidea | -0.218 | 0.628 | -0.347 | 0.729 |  |
| Salamandroidea | -0.759 | 0.660 | -1.152 | 0.249 |  |

Overall, Bd-exposed amphibians experienced higher mortality relative to controls (lnOR = 1.56 ±0.65 95% CI), but the magnitude of the effect of Bd exposure on Bd-related mortality varied depending on host life stage (Table 1 & Fig. 3). Hosts exposed to Bd as metamorphs experienced the highest odds of mortality (lnOR = 2.48 ±0.38 95% CI; *k* = 87), followed by adults (lnOR = 1.58 ±0.52 95% CI; *k* = 58), whereas larvae had the lowest odds of mortality (lnOR = 0.61 ±0.45 95% CI; *k* = 60). Additionally, we found a significant positive relationship between mortality and Bd dose (z = 4.00 *p* < 0.01; Table 1 & Fig. 3).

Finally, we found that some host taxa experienced significantly higher mortality from Bd than others (Table & Fig. 4). Bufonoidea had the highest mortality (lnOR = 3.25 ±1.18 95% CI; *k* =60) followed by Hyloidea (lnOR = 2.65 ±1.78 95% CI; *k* = 63) and then Ranoidea (lnOR = 1.34 ±1.23 95% CI; *k* = 47). Bd exposure did not significantly increase mortality for amphibians belonging to Salamandroidea (lnOR = 0.80 ±1.29 95% CI; *k* = 31). The number of effect sizes for Scaphiopodidae (*k* = 3) and Leiopelmatidae (*k* = 1) species were minimal so, we did not attempt to interpret those results.

**Figure 4.**
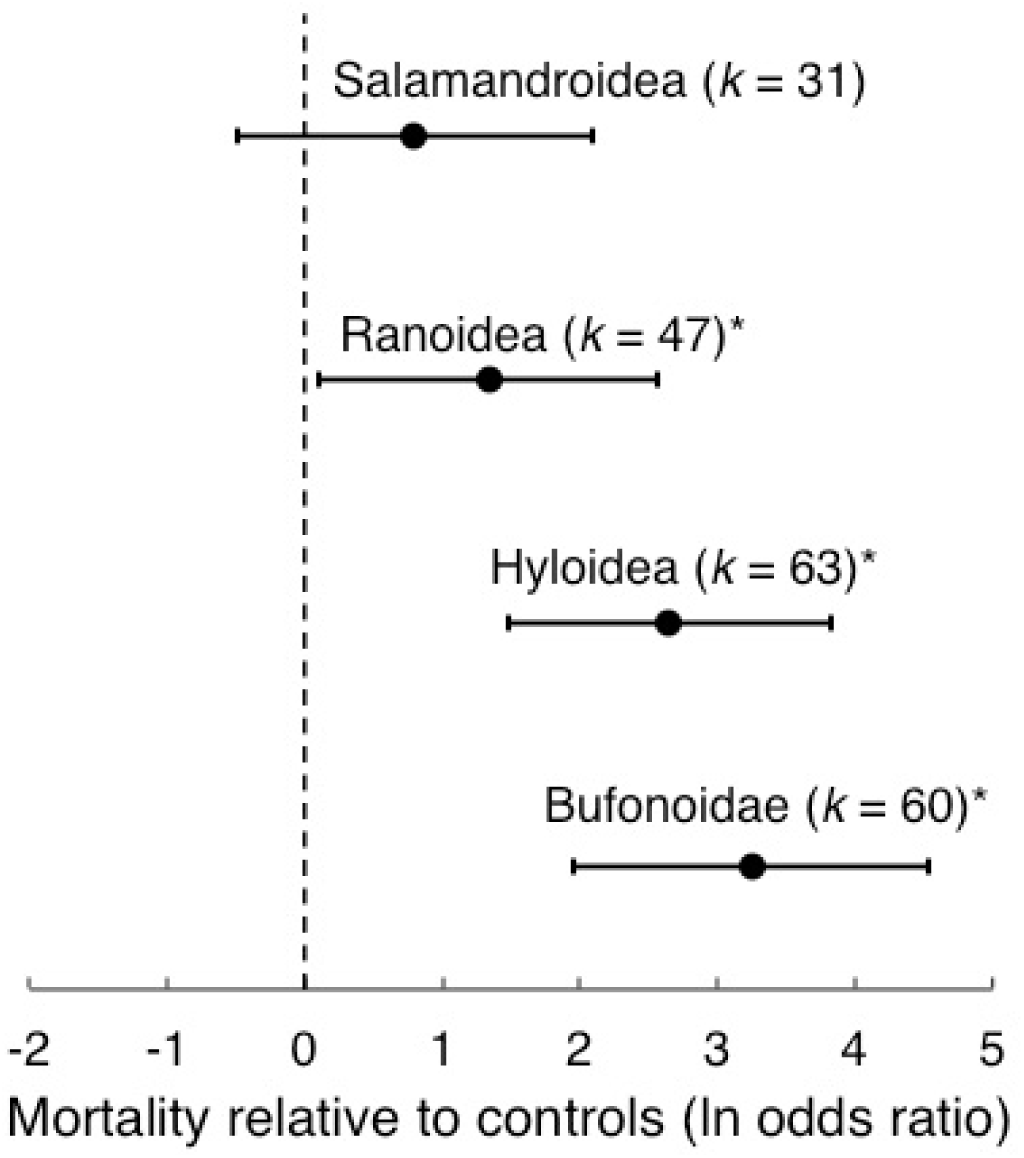
Forest plot of the marginal mean effect of Bd on host mortality relative to controls of taxonomic groups (presented as natural log transformed odds ratio). Positive values indicate greater mortality among Bd-exposed animals than among unexposed control animals. Asterisks indicate a significant difference from zero. Points represent the mean effect for that taxonomic group and error bars show associated 95% confidence intervals.

## Discussion

Species interactions can be sensitive to environmental conditions. Therefore, differences in laboratory and field conditions can reduce the transferability of empirical insights to management decisions. Here, we synthesized the effects of laboratory conditions that are easily manipulated by experimenters, such as temperature, study organism, developmental stage, and exposure dose. Our literature search highlighted a gap in laboratory studies of hosts and Bd isolates from the Middle East, Asia, and Africa (Fig. 1). Additionally, we found support for the thermal mismatch hypothesis: hosts from cooler climates were more susceptible to Bd at relatively warm lab temperatures, and vice versa (Table 1 & Fig. 2). Our data also show an overall positive effect of Bd exposure on mortality relative to controls (Table 1 & Fig. 3). In addition, we found that the strength of the effect of Bd exposure on mortality was dependent upon host life stage (Table 1 & Fig. 3) and host taxa (Table 1 & Fig. 4). Finally, we showed that Bd zoospore dose is positively related with mortality relative to controls, suggesting higher doses result in greater host mortality (Table 1 & Fig. 3).

Our literature search yielded a high number of effect sizes (*k* = 205) and included 47 amphibian species from 11 families as well as a 45 unique Bd isolates. However, we detected a geographic bias in the collection location of study organisms; the vast majority of hosts and Bd isolates were from North America and Europe. Less than 10% of our effect sizes represented host species collected from Central and South America or Oceania, and we did not find any effect sizes from Asia or Africa because no studies from these regions met our inclusion criteria. The distribution of amphibian study species matches up poorly with global amphibian diversity, which is higher in South America, Africa, and Asia than in North America or Europe (IUCN 2018). Because we were testing the thermal mismatch hypothesis, we did not include studies using hosts that were captive-bred beyond one generation in our study because: 1) hosts might have adapted to laboratory temperature, and 2) many studies lacked precise collection locations. This selection method excluded many studies from Oceania and Central America that used captive-bred imperiled or wild-extinct species out of necessity (Scheele et al. 2019). Furthermore, we were unable to find any studies that tested wild-collected or captive-bred Asian or African host species or Bd isolates that met our inclusion criteria. Future Bd research should consider laboratory experiments on species from these highly neglected areas, because of the strong genetic support for the recent emergence of Bd from northeastern Asia as well as the apparent lack of Bd-contributed declines in the region (O’Hanlon et al. 2018, Scheele et al. 2019).

Our meta-analysis supported the thermal mismatch hypothesis (Table 1 & Fig. 2). Cool-adapted hosts had the highest Bd-induced mortality at warm temperatures and warm-adapted hosts had the highest mortality at cool temperatures. Previous support for this hypothesis is based primarily on observations in the field and only three species studied in the laboratory (Cohen et al. 2017, Sauer et al. 2018, Cohen et al. 2019a, Cohen et al. 2019b), so our study demonstrates that these patterns also hold under controlled conditions for a broader range of species. Together, results from field and laboratory studies suggest that predicted increases in environmental temperatures caused by climate change might place cool-adapted species at greater risk of disease-related declines than warm-adapted species (Cohen et al. 2017, Sauer et al. 2018, Cohen et al. 2019a, Cohen et al. 2019b). Our results are unlikely to be driven by experiments that were purposely conducted at extreme temperatures because only five of the 58 studies included in this analysis manipulated environmental temperature. The vast majority of studies (49 of the 58 included in this analysis) simply conducted their experiments at a constant temperature without providing any justification for choosing that temperature. Researchers might be inadvertently impacting Bd growth and host mortality by conducting their experiments at temperatures that differ from the temperatures to which the host is adapted, potentially increasing host stress (Raffel et al. 2006). Outside of the mounting evidence for the thermal mismatch hypothesis in the Bd-amphibian system, there is a large body of research showing that environmental temperatures greatly impact experimental outcomes in this system as well as other amphibian-disease systems (Rojas et al. 2005, Paull et al. 2012, Brand et al. 2016). Researchers should carefully consider how experimental temperatures and host thermal preferences and tolerances impact the results and management applicability of conclusions drawn from laboratory experiments (Stevenson et al. 2014). Where possible, we encourage researchers to select temperatures that are ecologically relevant for their specific host-pathogen system.

As expected, we found an overall positive effect of Bd on host mortality relative to controls (Table 1 & Fig. 3) and the average minimum Bd dose needed to find an effect of Bd on host mortality varied across host life stage (Fig. 3). Amphibians exposed as metamorphs were more likely to experience mortality after Bd exposure than the larval or adult stages. Metamorphosis is energetically costly and there are likely trade-offs occurring between morphological development and the immunological function (Rollins-Smith 1998, Warne et al. 2011), which could make metamorphic amphibians more susceptible to Bd-induced mortality than larvae or adults. Larvae, while still susceptible to Bd, were less susceptible to Bd than metamorphs or adults (Table 1 & Fig. 3). Our review of the literature revealed that researchers tend to expose amphibians to similar doses of Bd, despite differences in susceptibility across life stages (mean log_10_ zoospore dose: larvae = 5.42 ± 0.21 SE; metamorphs = 5.07 ± 0.17 SE; adults = 5.95 ± 0.13 SE; Fig. 1b). Using lower Bd doses, when possible, may improve the ability of researchers to detect differences among treatment groups by preventing rapid death in all treatments due to heavy Bd infection (Carey et al. 2006). This is particularly true for metamorphic amphibians, which are on average dosed with 1000 times more zoospores than needed to find an effect on mortality (Fig. 1 & 3). Additionally, researchers interested in sub-lethal effects of Bd on amphibians might consider running experiments using larval or adult amphibians and/or using very low Bd zoospore doses (approximately < 10^2^ total zoospores; Fig. 1).

Finally, we found that Bufonoidea and Hyloidea species had the highest mortality risk, followed by Ranoidea species (Table 1 & Fig. 4). We were not able to detect a significant effect of Bd on Salamandroidea (Table 1 & Fig. 4), which supports field evidence that Salamandroidea species may be less susceptible to Bd than anurans (Bancroft et al. 2011). However, our analysis does not incorporate studies conducted with *Batrachochytrium salamandrivorans*, which has been associated with declines of salamander populations in Europe (Stegen et al. 2017). Our results are consistent with the severe risk of Bd-associated declines observed in tropical Bufonoidea and Hyloidea species (Scheele et al. 2019). Interestingly, all Bufonoidea species in our analysis are native to temperate regions and only one has been associated with declines, while most declining Bufonoidea are tropical. Thus, other factors such as local climate, thermal and hydric preferences, and small range sizes may be interacting with high host susceptibility to result in population declines in Bufonoidea. Conversely, there were many tropical Hyloidea species in our meta-analysis that have undergone Bd-associated declines or were collected from areas with severe Bd-associate declines.

Carefully designed experiments are especially important for understanding systems of conservation concern, including the amphibian-Bd system that has been associated with the decline of >500 amphibian species (Scheele et al. 2019). Additionally, our literature search highlighted the need for more research on hosts and Bd isolates from outside of Europe and North America. The regions with highest levels of amphibian declines (Scheele et al. 2019) and diversity (IUCN 2018) and the longest history of Bd (O’Hanlon et al. 2018) are some of the least studied regions in the world. Finally, our results are consistent with the thermal mismatch hypothesis, suggesting that there are context-dependent effects of environmental temperature on amphibian mortality in this system. This result highlights the need for researchers to carefully consider the thermal tolerances and optima of their host species before choosing an experimental temperature to avoid the confounding effect of thermal mismatch. In summary, because of their ability to alter experimental outcomes, our results suggest that factors such as experimental temperature and the life stages, densities, populations, and taxa of studied species should be carefully considered when designing species interaction experiments and subsequent interpretations for conservation and management.

## Supporting information

Metadata for Database S1

Database S1

## Acknowledgments

We thank O. Santiago for assistance assembling citations and standardizing nomenclature and L.C. Sackett and S. Deban for helpful comments on the manuscript. E.L.S. and J.R.R. conceived ideas, E.L.S., J.M.C., and T.A.M. oversaw construction of database, all authors extracted data from the literature to build the dataset, J.M.C. acquired climate data, E.L.S. and M.J.L. conducted statistical analyses, E.L.S. and J.R.R. wrote the first draft of the manuscript with comments from M.J.L., J.M.C., T.A.M., S.A.K., D.J.C., K.N., B.F.S. and E.A.R. Funds were provided by grants to J.R.R. from the National Science Foundation (EF-1241889, DEB-1518681), the National Institutes of Health (R01GM109499, R01TW010286-01), the US Department of Agriculture (2009-35102-0543), and the US Environmental Protection Agency (CAREER 83518801). Funding support was provided to D.J.C. and T.A.M. from the National Science Foundation (IOS-1755002 and ISO-1754862, respectively).

