## Supplementary material for "A meta-analysis reveals temperature, dose, life stage, and taxonomy influence host susceptibility to a fungal parasite": Metadata for Database S1

### Data S1

### Database S1: Database compiled from experimental studies that was used in the meta-analysis.

### Authors

Erin L. Sauer

Department of Integrative Biology, University of South Florida, Tampa, FL, 33620.

Department of Forest and Wildlife Ecology, University of Wisconsin, Madison, WI, 53706.

Jeremy M. Cohen

Department of Integrative Biology, University of South Florida, Tampa, FL, 33620.

Department of Forest and Wildlife Ecology, University of Wisconsin, Madison, WI, 53706.

Marc J. Lajeunesse

Department of Integrative Biology, University of South Florida, Tampa, FL, 33620.

Taegan A. McMahon

Department of Biology, University of Tampa, Tampa, FL, 33606.

David J. Civitello

Department of Biology, Emory University, Atlanta, GA, 30322.

Sarah A. Knutie

Department of Ecology and Evolutionary Biology, University of Connecticut, Storrs, CT, 06269.

Karena Nguyen

Department of Integrative Biology, University of South Florida, Tampa, FL, 33620.

Elizabeth A. Roznik

Department of Research and Conservation, Memphis Zoo, Memphis, TN, 38112.

Brittany F. Sears

BioScience Writers, Houston, TX, 77098.

Scott Bessler

Department of Integrative Biology, University of South Florida, Tampa, FL, 33620.

Bryan K. Delius

Department of Integrative Biology, University of South Florida, Tampa, FL, 33620.

Neal Halstead

Wildlands Conservation, Tampa, FL, 33647.

Nicole Ortega

Department of Integrative Biology, University of South Florida, Tampa, FL, 33620.

Matthew D. Venesky

Department of Biology, Allegheny College, Meadville, PA, 16335.

Suzanne Young

Environmental Engineering Institute, Ecole polytechnique fédérale de Lausanne (EPFL), Lausanne, Switzerland.

Jason R. Rohr

Department of Integrative Biology, University of South Florida, Tampa, FL, 33620.

Department of Biological Science, University of Notre Dame, South Bend, IN, 46556.

### File list (files found within DataS1.zip)

DatabaseS1.csv

**Description**

DatabaseS1.csv – Description of column contents in format: Column header – Description (unit, if applicable).

Paper - Citation

Order - Host order

Family	- Host family

Superfamily - Host taxonomic group (see Table S1)

Genus - Host genus

Species - Host species

Host.Lat - Latitude of the host collection site (decimal degrees)

Host.Long - Longitude of the host collection site (decimal degrees)

Host.BIO1 - 50-year mean temperature at the location from which the host was collected (degrees C)

Lifestage - Life stage of the host when exposed to Bd

N.Treatment - Treatment group sample size

Mortality.Treatment - Mortality in the treatment group

Dt - Number of animals dead in the treatment group at the end of the experiment

At - Number of animals alive in the treatment group at the end of the experiment

N.Control - Control group sample size

Mortality.Control - Mortality in the control group

Dc - Number of animals dead in the control group at the end of the experiment

Ac - Number of animals alive in the control group at the end of the experiment

ln.or - Natural log odds ratio (see Eqn. 1)

v.ln.or - Variance of lnOR (see Eqn. 2)

Study.Treatment - Treatment group name given in the study

Temp - Experimental temeperature (degrees C)

Bd.Strain - Bd isolate used in the study

Bd.Lat - Latitude of the Bd collection site

Bd.Long - Longitude of the Bd collection site

Dose - Bd dose (number of Bd zoospores)

logDose - Log10 (zoospore dose)

Duration - Duration of the experiment (days)
