## Supplemental information for "A meta-analysis reveals temperature, dose, life stage, and taxonomy influence host susceptibility to a fungal parasite"

Appendix S1 for: Erin L. Sauer, Jeremy M. Cohen, Marc J. Lajeunesse, Taegan A. McMahon, David J. Civitello, Sarah A. Knutie, Karena Nguyen, Elizabeth A. Roznik, Brittany F. Sears, Scott Bessler, Bryan K. Delius, Neal Halstead, Nicole Ortega, Matthew D. Venesky, Suzanne Young, and Jason R. Rohr. 2019. A meta-analysis reveals temperature, dose, life stage, and taxonomy influence host susceptibility to a fungal parasite. *Ecology*.

**Supplemental methods**

*Data collection*

Search results were screened using the *abstract_screener* function of the *metagear* package [1] (final count: 58 studies; [2-59]). We manually extracted data from text and tables, and extracted data from figures using Plot Digitizer version 2.6.6 (plotdigitizer.sourceforge.net). In addition, we recorded other information about the biology and methodology of each study, including (if available): host species, geographic coordinates where hosts were collected, developmental stage (larvae, metamorph, or adult), percent mortality in treatment and control animals, sample sizes, Bd isolate, geographic coordinates of the collection site of the isolate, Bd dose (total zoospores), and duration of the experiment in days. Species nomenclature was standardized according to the IUCN [60]. Finally, for each row in our dataset, we extracted annual mean temperature, mean annual minimum temperature, and mean annul maximum temperature across 50 years (1950-2000) from WorldClim (www.worldclim.org, *raster* package, *extract* function in R 3.5.1 [61]) that corresponded to the location where each host population was collected.

We extracted climatic variables to estimate host-adapted temperature, which is needed to test for a thermal mismatch effect. This effect is represented by an interaction between host-adapted temperature and the laboratory temperature at which the experiment was conducted (Temp). Ultimately, we included 50-year mean temperature (Tmean) at the host’s collection site as the assumed host-adapted temperature. Mean annual minimum (Tmin) and maximum temperatures (Tmax) yield very similar results (Temp * Tmean: β = -0.02, *z* = 2.75, *p* < 0.01; Temp * Tmin: β = -0.02, *z* = 2.77, *p* < 0.01; Temp * Tmax: β = -0.02, *z* = 2.38, *p* = 0.02). This is not surprising as mean annual minimum and maximum temperature are highly colinear with each other and with 50-year mean temperature.

Without laboratory measures of peak performance temperature for every population of hosts in the meta-analysis, 50-year mean temperature from the host’s collection location is likely the best estimate of host-adapted temperature. However, the 50-year mean temperature from the host’s collection location does not need to perfectly match the host’s peak performance, as long as the host’s thermal performance breadth is roughly centered (asymmetrically) on the host’s peak performance temperature. Thermal mismatch models suggest that as hosts encounter increasingly dissimilar temperatures, disease risk goes up continuously, because the models are linear. Thus, if the temperature is only a few degrees different than the host’s adapted temperature, the increase in risk is small, but if it is 20 degrees off, the increase in risk is large. The nature of this linear increase in risk as the magnitude of the thermal mismatch increases allows our estimate of peak host performance temperature to be approximate.

*Taxonomic consolidation*

In order to explore differences in susceptibility among host taxa, species were consolidated into taxonomic groups with larger sample sizes. Thus, taxonomic groups represent either a superfamily (Bufonoidea, Hyloidea, Ranoidea, and Pelobatoidea), or a suborder (Salamandroidea and Archaeobatrachia). We were unable to further consolidate Pelobatoidea (k = 3) or Archaeobatrachia (k = 1) species as they are distantly related to the other groups and each other, thus, we do not present results or attempt to interpret the effect of Bd on those two groups with very small sample sizes (see Table S1 for full list of host species included in the meta-analysis).

*Figure generation*

We created partial residual plots to visualize the main effects and interactions in our model (Figs. 2, 3, & 4). Partial residual plots show a predicted relationship between a predictor and response variable while controlling for the other variables in the model [62]. To make the plots, we first built an identical model using a Bayesian linear mixed-effects model (*blme* package, *blmer* function). In this model, each effect size was weighted by the inverse of that effect size’s variance and the residual variance prior was fixed at one, meaning the weights of each effect size were assumed to be exactly known [63] (see Table S2 for model coefficient and standard error comparisons). We used this approach because visualization tools for mixed-effects meta-analytic models in *metafor* are currently limited. Finally, we generated partial residual plots (*visreg* package, *visreg* function [62]) from the blmer model to visualize the main effects of experimental duration (Fig. 3) and dose (Fig. 4) as well as the interaction between laboratory and long term temperature (Fig. 2).

| Table S1\| Taxonomic grouping of all species included in DataS1: Database S1. | | | | |
| --- | --- | --- | --- | --- |
| Superfamily or suborder | Family | Genus | Species | *k* |
| Bufonoidea | Bufonidae | Anaxyrus | *Anaxyrus americanus* | 5 |
|  |  |  | *Anaxyrus boreas* | 24 |
|  |  |  | *Anaxyrus fowleri* | 2 |
|  |  |  | *Anaxyrus terrestris* | 1 |
|  |  | Bufo | *Bufo bufo* | 21 |
|  |  |  | *Bufo quercicus* | 5 |
|  |  | Incilius | *Incilius nebulifer* | 2 |
| Hyloidea | Brachycephalidae | Brachycephalus | *Brachycephalus pitanga* | 1 |
|  |  | Ischnocnema | *Ischnocnema parva* | 1 |
|  | Craugastoridae | Craugastor | *Craugastor fitzingeri* | 1 |
|  | Hylidae | Agalychnis | *Agalychnis callidryas* | 1 |
|  |  | Dendropsophus | *Dendropsophus meridensis* | 1 |
|  |  |  | *Dendropsophus minutus* | 1 |
|  |  | Hyla | *Hyla chrysoscelis* | 7 |
|  |  |  | *Hyla versicolor* | 8 |
|  |  | Litoria | *Litoria ewingii* | 1 |
|  |  |  | *Litoria raniformis* | 1 |
|  |  |  | *Litoria verreauxii* | 3 |
|  |  | Osteopilus | *Osteopilus septentrionalis* | 15 |
|  |  | Pseudacris | *Pseudacris crucifer* | 2 |
|  |  |  | *Pseudacris feriarum* | 1 |
|  |  |  | *Pseudacris regilla* | 13 |
|  |  |  | *Pseudacris triseriata* | 3 |
|  | Leptodactylidae | Eleutherodactylus | *Eleutherodactylus coqui* | 3 |
| Archaeobatrachia | Leiopelmatidae | Leiopelma | *Leiopelma hochstetteri* | 1 |
| Ranoidea | Ranidae | Lithobates | *Lithobates catesbeianus* | 7 |
|  |  |  | *Lithobates clamitans* | 2 |
|  |  |  | *Lithobates pipiens* | 4 |
|  |  |  | *Lithobates sphenocephalus* | 2 |
|  |  |  | *Lithobates sylvaticus* | 7 |
|  |  | Rana | *Rana cascadae* | 13 |
|  |  |  | *Rana muscosa* | 4 |
|  |  |  | *Rana pretiosa* | 2 |
|  |  |  | *Rana subaquavocalis* | 1 |
|  |  |  | *Rana temporaria* | 3 |
|  |  |  | *Rana yavapaiensis* | 2 |
| Salamandroidea | Ambystomatidae | Ambystoma | *Ambystoma californiense* | 2 |
|  |  |  | *Ambystoma laterale* | 2 |
|  |  |  | *Ambystoma tigrinum* | 1 |
|  | Plethodontidae | Desmognathus | *Desmognathus monticola* | 2 |
|  |  |  | *Desmognathus orestes* | 1 |
|  |  | Plethodon | *Plethodon cinereus* | 9 |
|  |  |  | *Plethodon glutinosus* | 3 |
|  |  |  | *Plethodon metcalfi* | 2 |
|  | Salamandridae | Lissotriton | *Lissotriton helveticus* | 3 |
|  |  | Notophthalmus | *Notophthalmus viridescens* | 6 |
| Pelobatoidea | Scaphiopodidae | Scaphiopus | *Scaphiopus holbrookii* | 3 |

| Table S2\| Comparison of the Bayesian linear mixed-effects model *(blme* model) used to make Figures 2, 3, & 4 and the linear mixed-effects model described in the main text and in Table 1 *(metafor* model). The main effect of Bd on mortality is indicated by the grand mean where the coefficient represents the pooled effect of Bd on mortality as a log odds ratio. For continuous variables, the coefficient indicates the direction of the effect. For categorical variables (life stage & taxonomic group only), the sign of the coefficient indicating the direction of the difference from the grand mean. | | | | |
| --- | --- | --- | --- | --- |
|  | *blme* model | | *metafor* model | |
|  | Coefficient | SE | Coefficient | SE |
| Grand mean | 1.559 | 0.314 | 1.559 | 0.314 |
| LongTermTemp | 0.288 | 0.133 | 0.288 | 0.133 |
| LabTemp | 0.189 | 0.094 | 0.189 | 0.094 |
| LongTermTemp*LabTemp | -0.019 | 0.007 | -0.019 | 0.007 |
| Duration | 0.009 | 0.004 | 0.009 | 0.004 |
| Larvae | -0.947 | 0.228 | -0.947 | 0.228 |
| Metamorph | 0.924 | 0.195 | 0.924 | 0.195 |
| Adult | 0.023 | 0.266 | 0.023 | 0.266 |
| logDose | 0.454 | 0.114 | 0.454 | 0.114 |
| Bufonoidea | 1.690 | 0.657 | 1.690 | 0.657 |
| Hyloidea | 1.093 | 0.601 | 1.093 | 0.601 |
| Ranoidea | -0.218 | 0.628 | -0.218 | 0.628 |
| Salamandroidea | -0.759 | 0.660 | -0.759 | 0.660 |

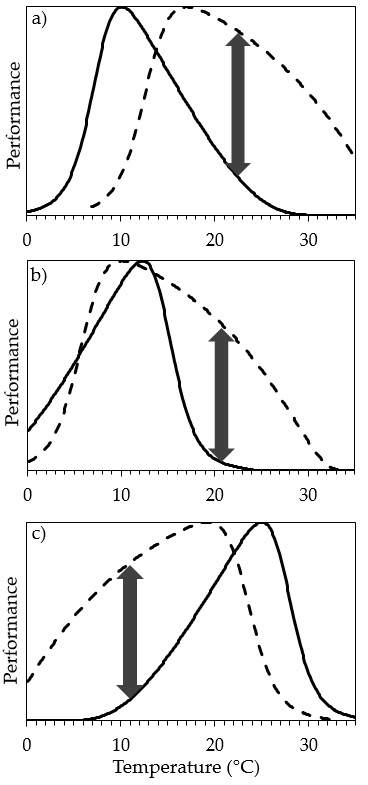

**Figure S1. Predictions of the *thermal mismatch hypothesis* are robust to underlying assumptions [reprinted from 64].** The *thermal mismatch hypothesis,* which predicts that hosts should be more susceptible to outbreaks when environmental conditions shift away from their optima, is based on a few key underlying assumptions: 1) hosts and parasites are locally adapted to their thermal environments, 2) cold- and warm-adapted hosts and parasites have right- and left-skewed thermal performance curves, respectively, and 3) parasites have broader thermal breadths than hosts. However, the predictions that arise from this hypothesis, that cold- and warm-adapted hosts and parasites should have outbreaks at relatively warm and cold temperatures, respectively, are robust to several of these assumptions. **a)** Even when parasites (dashed line) and hosts (solid line) do not have similar performance peaks as might be expected with local adaptation (as may be the case with *Bd*), cold- and warm-adapted hosts still often have outbreaks at relatively warm and cold temperatures, respectively. **b)** Likewise, peak growth of parasites can occur at warm temperatures on cold-adapted hosts regardless of whether their performance curves are right- or left-skewed. **c)** In situations where host thermal breadth eclipses parasite thermal breadth on the right (warm) end of the curve, peak parasite growth is likely to occur at cold temperatures. This scenario may often occur in the amphibian/*Bd* system. Hence, the predictions of the thermal mismatch hypothesis are robust to several of the assumptions. See Fig. 1 to compare the results in this figure to those where the results are restricted by the three assumptions above.
